# Bayesian Variable Selection Approaches to Identifying Selection in Evolutionary Datasets

**DOI:** 10.64898/2026.10.19.760252

**Authors:** Yongkang Li, Scott C. Schmidler

## Abstract

We consider the problem of identifying individual sites and characters under selection in phylogenetic datasets. We introduce an approach based on the application of Bayesian variable selection to mutation-selection models, which enables both sites and selection coefficients to be explicitly excluded from the model in order to control parameter complexity. We provide an accelerated Gibbs sampling algorithm for model selection and parameter estimation, which enables our approach to scale to realistic sequence lengths and dataset sizes. We compare this approach to previous models that do not produce sparse solutions but attempt to control complexity through Dirichlet-process (DP) mixture model prior distributions. We demonstrate that the Bayesian variable selection approach outperforms the DP approach for a wide range of settings and selection signal strengths, while offering additional advantages of sparsity and interpretability. We also identify a problem of the DP prior producing “phantom selection” artifacts, on both simulated and real datasets, which the Bayesian variable selection approach avoids.

## 1 Introduction

Identifying sites under selection is a central problem in molecular evolution. Classical approaches infer selection using the ratio *dN/dS* of nonsynonymous to synonymous substitution rates under a codon substitution model (Goldman and Yang 1994; Muse and Gaut 1994). Extensions allowing *dN/dS* to vary across sites enable the detection of site-specific selection (Nielsen and Yang 1998; Yang et al. 2000, 2005; Kosakovsky Pond and Frost 2005; Huelsenbeck et al. 2006; Murrell et al. 2012, 2013). However, these methods do not directly estimate the selective preferences for individual amino acids. Moreover, elevated dN/dS need not imply positive selection on amino acids when synonymous codons differ in fitness (Spielman and Wilke 2015). For sequences sampled within a population, *dN/dS* may also be nonmonotonically related to selection strength, complicating its interpretation (Kryazhimskiy and Plotkin 2008).

Mutation-selection models instead derive codon substitution rates from mutation rates and fixation probabilities determined by fitness differences (Halpern and Bruno 1998; Sella and Hirsh 2005; Yang and Nielsen 2008). Site-specific amino-acid fitnesses are represented by normalized propensity vectors, permitting direct estimation of relative amino-acid preferences (Rodrigue et al. 2010; Tamuri et al. 2012, 2014). These models have also been extended to detect adaptive evolution (Rodrigue and Lartillot 2017; Rodrigue et al. 2021), and to incorporate experimentally measured amino-acid preferences (Bloom 2014, 2017). Failure to account for such site-specific amino-acid preferences can also lead to biased estimates of branch lengths and tree topology during phylogenetic inference (Lartillot et al. 2007; Castoe et al. 2009; Hilton and Bloom 2018).

A major challenge of identifying selection with site-heterogeneous mutation-selection models is the high dimensionality: each site requires 19 free parameters to describe the relative fitnesses of 20 amino acids. When sequence divergence or sample size is limited, these parameters may be weakly identified. Existing approaches control model complexity through penalized likelihood (Tamuri et al. 2014), Bayesian approaches using hierarchical priors that shrink site-specific fitness profiles toward a shared population distribution (Rodrigue 2013), or more sophisticated nonparametric Bayes approaches based on Dirichlet-process (DP) mixtures that cluster sites into shared fitness profiles (Lartillot and Philippe 2004; Rodrigue et al. 2010; Rodrigue and Lartillot 2014). One shortcoming of these approaches from a user’s perspective is that they generally do not provide an explicit estimate of the probability of the presence or absence of site- or amino-acid-specific fitness effects, so identifying selection requires an additional decision rule or thresholding procedure. Moreover, as we will demonstrate below, pooling fitness profiles across sites can assign strong amino-acid preferences to weakly informative sites based primarily on evidence from other sites. Such estimates can produce what we term *phantom selection*, in which the inferred site preferences correspond to amino acids never observed in the dataset.

We address these limitations by formulating selection identification as a hierarchical Bayesian variable selection (BVS) problem (Mitchell and Beauchamp 1988; George and McCulloch 1993). Hierarchical point-mass priors enable the exclusion of site-specific fitness differences and individual amino-acid coefficients, providing posterior inclusion probabilities separately from estimates of effect magnitude. To overcome the slow mixing of conventional Gibbs samplers under this hierarchy, we employ an ancillarity-sufficiency interweaving strategy (ASIS) (Yu and Meng 2011; Li and Schmidler 2026) that combines complementary parameterizations within each MCMC iteration.

We evaluate the sensitivity and specificity of BVS across datasets with varying selection signal strengths, and compare the resulting estimation accuracy with that obtained using state-ofthe-art non-parametric Bayesian priors using DP-mixtures. We further analyze three datasets from the literature: alignments of vertebrate *β*-globins, succinate dehydrogenase subunit B (SDHB), and class A *β*-lactamases. For SDHB, we examine correspondence between inferred amino-acid preferences and experimentally characterized functional residues. For *β*-lactamase, deep mutational scanning measurements (Stiffler et al. 2015) provide an independent experimental reference. Across these analyses, BVS avoids the phantom selections observed under the DP mixture and, for *β*-lactamase, its inferred most-preferred amino acids at highly ranked sites show greater agreement with experimental measurements.

The remainder of this paper is organized as follows. Section 2 reviews phylogenetic mutation-selection models and their parameter complexity, and briefly reviews the DP-mixture approach. Section 3 introduces the hierarchical BVS model and Section 4 describes the accelerated MCMC algorithm. Section 5 describes empirical results on simulated and real datasets.

## 2 Background

### 2.1 Molecular Evolution Models and Phylogenetic Inference

Modern molecular phylogenetic inference is commonly performed using maximum-likelihood or Bayesian methods under explicit stochastic models of molecular sequence evolution. Standard evolutionary models assume that each site (nucleotide, codon, or amino acid) in a sequence evolves independently according to a continuous-time Markov chain (CTMC) specified by a time-homogeneous rate matrix **Q**^(*i*)^ with rates 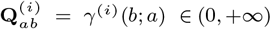 specifying the rate at which the site *i* mutates from *a* to *b*. Sequence transition probabilities can then be written as

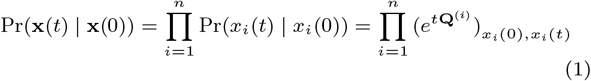

where **x**(*t*) = (*x*_1_(*t*), … , *x*_*n*_(*t*)) denotes the state of the sequence at time *t*. Given a set of observed sequences **X** = (**x**^1^, … , **x**^*m*^) at the tips of a tree *g* (including topology and branch lengths), the likelihood of each individual site can be computed efficiently using Felsenstein’s pruning algorithm (Felsenstein 1981), and the full sequence likelihood obtained as the product of site likelihoods (1). Maximizing the full sequence likelihood over tree topology and branch lengths yields maximum-likelihood tree estimates (Guindon and Gascuel 2003; Stamatakis 2014), while incorporation of priors and application of Markov chain Monte Carlo (MCMC) sampling techniques yields (samples from) the posterior distribution over trees (Huelsenbeck and Ronquist 2001; Drummond and Rambaut 2007). Similar calculations enable posterior sampling of complete substitution histories along each branch, providing a full stochastic reconstruction of ancestral states and substitution events conditional on the observed sequences. However, such model-based analyses are premised on the assumed substitution model **Q**: for example, if the true evolutionary process is shaped by site-specific selection at the amino acid level – and the substitution model fails to account for this – the analysis can systematically mis-estimate branch lengths, ancestral states, and tree topology (Castoe et al. 2009; Hilton and Bloom 2018). Hence, accounting for selection explicitly in the substitution model is often essential for accurate phylogenetic inference.

### 2.2 Mutation-Selection Models

Significant progress has been made in incorporating selection into phylogenetic modeling. Selection has traditionally been tested for using nonsynonymous-to-synonymous rate ratios (Goldman and Yang 1994; Muse and Gaut 1994), with extensions to site heterogeneous ratios enabling detection of specific sites under selection (Nielsen and Yang 1998; Yang et al. 2000). More recently, *mutation-selection models* explicitly link substitution rates to site-specific amino-acid fitness, enabling direct estimation of selection strength from sequence data (Halpern and Bruno 1998; Yang et al. 2000; Yang and Nielsen 2008; Teufel et al. 2018). These models often retain the independent-and-identically-distributed (or exchangeable) mutation process across sites, while allowing fitness profiles to vary across sites. These models have been used within Bayesian and maximum-likelihood frameworks to infer fully site-heterogeneous fitness profiles (Rodrigue et al. 2010, 2021).

To account for fitness at the amino acid level, selection coefficients are incorporated into a codon mutation model (Goldman and Yang 1994; Muse and Gaut 1994; Nielsen and Yang 1998) as follows. Let C be the set of 61 sense codons, and A the set of 20 amino acids. A protein-coding nucleotide sequence **x** = (*x*_1_, *x*_2_, … , *x*_*l*_) of length *l* = 3*I* is modeled as a sequence of *I* codons, (*c*_1_, … , *c*_*I*_), where *c*_*i*_ = (*x*_3*i−*2_, *x*_3*i−*1_, *x*_3*i*_) ∈ C ⊂ N^3^ for each 1 ≤ *i* ≤ *I*. Let *φ* : C → A be the function mapping a codon to its encoded amino acid. The mutation-selection model (Halpern and Bruno 1998; Yang et al. 2000; Lartillot 2006; Yang and Nielsen 2008) accounts for selection acting on the amino acid sequence, with the fitness of amino acids allowed to vary by site. For each amino acid site *i* (corresponding to codon *c*_*i*_), define a vector of amino acid propensities, **Ψ**_*i*_ = (Ψ_*ij*_)_*j∈A*_, with Ψ_*ij*_ *>* 0 and _*j∈A*_ Ψ_*ij*_ = 1. The evolution of each codon *c*_*i*_ is then described by a distinct 61 × 61 rate matrix **Q**^(*i*)^ with rate of substitution from codon *c* to codon *c*^*′*^ at site *i* given by:

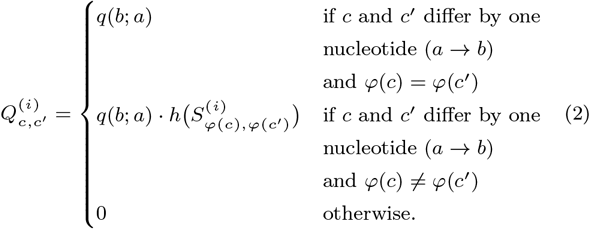

Here *q*(*b*; *a*) is the nucleotide substitution rate from a 4 × 4 nucleotide rate matrix **Q** = (*q*(*b*; *a*)); the scaled selection coefficient 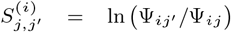 represents the log-fitness difference between amino acids *j* and *j*^*′*^ at site *i* and can equivalently be expressed as 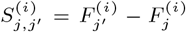, where 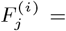 ln Ψ_*ij*_ denotes the individual amino-acid log-fitness; 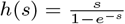 is the fixation factor for a mutation with scaled selection coefficient *s*, defined as the ratio of the fixation probability under selection to that under neutrality. The multiplicative form (2) follows from the weak-mutation approximation (Golding and Felsenstein 1990). Under this model, the stationary distribution of codons at site *i*, denoted as ***ζ***_*i*_ = (*ζ*_*i,c*_)_*c*_*∈C*, is given by:

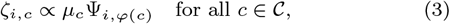

where *µ*_*c*_ are the stationary codon frequencies obtained under nucleotide mutation model **Q**. This model captures site interdependencies within individual codons induced by selective pressure, while retaining the simplicity of the independent site assumption at the level of both nucleotide mutations and codon sites.

### 2.3 Model Complexity

The site-heterogeneous fitness formulation substantially increases the model dimension relative to standard mutation models, with the amino-acid propensity vector **Ψ**_*i*_ contributing 19 additional free parameters for each site. When the number of observed sequences is small or sequence divergence is limited, this may exceed the amount of information available in the data for estimating parameters. An example is given by the globin dataset considered in Section 5, where a handful of leaf sequences must inform a fitness profile at every codon site. As a result of this over-parameterization, site-specific fitness profiles estimated with limited data are prone to overfitting and high variance, with 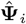 at risk of fitting noise in the data rather than genuine selective pressure, due to relatively few substitutions being observed at a given site.

In addition, scientists often desire parsimonious models to enhance understanding. Many or even most codon sites in typical coding sequences may not be under strong site-specific selection due to the substantial degeneracy in the sequence-structure mapping. Thus the inclusion of 19 free selection parameters per site can introduce unnecessary model complexity for the majority of sites.

This potential over-parameterization is well recognized in the field, and some mitigating solutions have been proposed. In some cases, fitness profiles can be derived from experimental measurements such as amino acid preferences measured directly via deep mutational scanning (Bloom 2014). However, such exhaustive experimental data are not available for most problems, and fitness profiles are instead estimated directly using the observed sequences. In this situation, Bayesian methods can provide a principled mechanism for controlling model complexity through the use of hierarchical or structured prior distributions on the set of fitness profiles 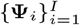. The CAT model (Lartillot and Philippe 2004) and its mutation-selection extensions (Rodrigue et al. 2010; Rodrigue and Lartillot 2014) use Dirichlet process mixture (Ferguson 1973) priors to cluster sites into a smaller number of shared fitness categories, with

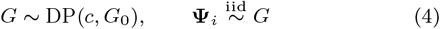

where *c* is a concentration parameter and *G*_0_ a base distribution defined on the probability simplex of amino-acid propensity vectors, while related constructions parameterize the base distribution *G*_0_ of the DP by a shared mean profile and a concentration parameter controlling variation around that mean (e.g. Dirichlet(*κ****ξ***) (Lartillot et al. 2007; Rodrigue 2013)). Such *shrinkage* priors reduce the effective number of free parameters by pooling information across sites: when the data at an individual site are limited, posterior estimates are pulled toward the shared population distribution, while sites at which the data are sufficiently informative are allowed to deviate.

## 3 Bayesian Variable Selection for Mutation-Selection Models

While shrinkage methods provide a popular solution to controlling model complexity and overfitting in high-dimensional parameter estimation, they are not the only solution. An alternative is the use of model selection or hypothesis testing methods to remove unneeded parameters from the model entirely. In many domains where parsimony and interpretability of the resulting model is valued as much as pure predictive performance, these methods offer an attractive alternative. Under the Bayesian framework, one such approach is the use of *Bayesian variable selection* (Mitchell and Beauchamp 1988; George and McCulloch 1993, 1997). Here we consider the use of Bayesian variable selection-style techniques to control the complexity of site-heterogeneous mutation-selection models in a rigorous but parsimonious way, enabling the model to identify and capture selective pressure where it exists, while removing parameters completely where evidence for selection is lacking in the data.

### 3.1 Controlling Complexity Using Model-Selection Priors

In contrast to the shrinkage estimation described in Section 2.3, model selection approaches control model complexity by explicitly removing some parameters from the model. In the Bayesian context, this is often performed by the use of model-selection priors, which encourage parameters to drop out of the model completely in the absence of evidence in the data that they are needed. Rather than merely shrinking weak effects toward zero, model-selection priors assign positive prior probability to the lower dimensional subspace obtained by setting some parameters exactly equal to zero. (This distinction parallels the contrast in classical analyses between *L*_2_*- (and L*_1_*-) regularization* in regression, where an *L*_2_-norm (ridge) penalty shrinks all coefficients toward zero but leaves them nonzero, and *subset selection* methods (Miller 1990) which add and remove variables from the regression equation completely. *L*_1_-norm (lasso) penalties achieve some of both effects, shrinking some coefficients while driving others to exactly zero, yielding a potentially sparse subset (Hoerl and Kennard 1970; Tibshirani 1996; Ishwaran and Rao 2005; Bai et al. 2021)).

The canonical example of a model selection prior is the pointmass prior, constructed as a mixture distribution which assigns a positive probability of a parameter *θ* taking exactly the value zero (Mitchell and Beauchamp 1988):

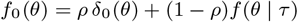

for some continuous distribution *f* (· | *τ*). One advantage of this approach is that the resulting posterior distribution provides both the estimated magnitude of an effect and a *posterior inclusion probability* which gives an explicit estimate of the probability that the effect exists at all. Point-mass priors also possess several attractive theoretical properties (George and McCulloch 1993; Ishwaran and Rao 2005; Castillo et al. 2015; Bhadra et al. 2019) including consistent recovery of the true model in the limit, while the marginalization over models to compute posterior inclusion probabilities represents a form of Bayesian *model averaging* (Kass and Raftery 1995; Hoeting et al. 1999), which appropriately propagates uncertainty in the model structure rather than conditioning inference on a single selected model. More recent global-local priors (Carvalho et al. 2010; Bhattacharya et al. 2015; Piironen and Vehtari 2017) provide a continuous alternative, combining substantial concentration near zero with heavy tails that permit strongly supported signals to escape shrinkage, and can have computational advantages. Extension of global-local priors to hierarchical group inclusion structures such as the site- and amino-acid-levels here is less clear; as a result we consider only point-mass priors here.

Point-mass priors also have the advantage of separating inference on the *presence* of a selection effect and its estimated *magnitude*: a site with a large estimated selection coefficient but low inclusion probability signals that neutrality cannot be confidently ruled out, but the selection effect – if present – may be large. This distinction is not captured directly by DP-mixture priors or continuous shrinkage priors, where true sparseness is only achieved by imposing thresholding values, and the resulting inclusion probabilities may be highly sensitive to the choice of threshold. Thus model-selection priors are particularly well suited for mutation-selection models, where parsimony and interpretability of results are of substantial interest, and the presence or absence of selection effects often of significant interest in its own right. Bayesian variable selection approaches provide these benefits while retaining full flexibility for sites where the evidence indicates selection.

### 3.2 Point-mass Priors for Selection Coefficients

Here we treat selection coefficient inclusion as a Bayesian variable selection (BVS) problem. Our approach uses a hierarchical representation for the presence of selection that enables selection to be included or excluded at both the site- and amino-acid-specific levels. We adopt the use of point-mass mixture priors for each site’s profile consisting of a mixture of a point-mass (an atom of positive probability mass) at exactly the neutral value 0 with a continuous “slab” for non-zero selection coefficients. We will see that pointmass priors of this form provide direct, interpretable answers to two key questions in our application: which codon sites and amino acids show evidence of selection, and, the magnitude and direction of that selection.

#### Site selection

We assume that each codon site is either *fully neutral* or *under selection* (positive or negative), indicated by a latent binary vector ***z*** = (*z*_1_, … , *z*_*I*_), where *z*_*i*_ ∈ {0, 1} for all 1 ≤ *i* ≤ *I*. Each latent variable *z*_*i*_ is assigned a Bernoulli prior:

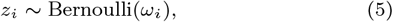

with *ω*_*i*_ the prior probability assigned to the hypothesis *under selection* and 1−*ω*_*i*_ the prior probability assigned to the hypothesis *neutral*. In some applications, informative *ω*_*i*_’s may be obtained by knowledge of the problem, such as structural information, sequence alignment databases, or knowledge of functional locations; for example the framework vs. complementarity determining (CDR) regions of antibodies. Here we do not assume such prior information is available; instead we allow the model to infer these site-specific probabilities by use of a hierarchical prior. That is, the *ω*_*i*_’s are assumed to follow a Beta hyperprior:

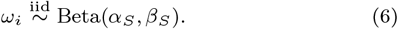

#### Neutral sites

If *z*_*i*_ = 0, the site is considered fully neutral. All amino acids are considered to have equal fitness, and the conditional prior distribution over propensity vectors is a point mass at the uniform distribution:

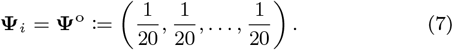

#### Amino acid selection and fitness profiles

If *z*_*i*_ = 1, the site is governed by a model that allows for positive, neutral, and negative amino-acid-specific selection. This is parameterized using a second level of inclusion indicators (*z*_*ij*_)_*j∈A*_, and corresponding hyperprior, for site- and amino-acid-specific selection intensities *η*_*ij*_ ∈ R:

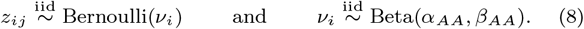

These hyperparameters are typically chosen to favor neutrality (*α*_S_ ≪ *β*_S_, *α*_AA_ ≪ *β*_AA_) to encourage sparsity. Conditional on the site-and-amino-acid selection coefficient being included in the model, the prior for its magnitude is specified by an exchangeable logistic-normal distribution on the simplex, with 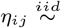 Normal(0, *σ*^2^) and the propensity vector **Ψ**_*i*_ is constructed as

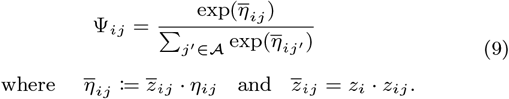

where 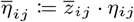 and 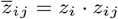.

The definition of fitness coefficients in relative terms (9) creates an inherent non-identifiability, which we resolve by imposing a site-specific centering constraint ∑_*j*_ *η*_*ij*_ = 0 for every *i* to remove invariance under additive shifts.

The formulation (5-9) allows the model to infer both which sites are under selection, and which amino acids are under selection (and in which direction, positive or negative) at a given codon site, providing a powerful yet parsimonious description of site-specific evolution. The choice of the hyperparameters *α*_S_, *β*_S_, *α*_AA_, *β*_AA_ reflects the level of sparsity desired in moderately informative datasets, allowing the BVS priors to emphasize sites and amino acids with the strongest evidence, thus providing a ranked, interpretable set of the most evidence-supported instances of selection compared to the dense profile obtained with standard mutation-selection models, in which genuine signal can be difficult to separate from noise. The appropriate degree of sparsity can be set according to the amount of data available: with a large alignment or a well-resolved tree, *α*_S_, *α*_AA_ can be set large relative to *β*_S_, *β*_AA_, placing more prior mass on inclusion without inflating the false-positive rate; while with limited data, or when conservative reporting is preferred, *α*_S_, *α*_AA_ can be set small relative to *β*_S_, *β*_AA_ so that only sites and amino acids with strong support will be reported. Meanwhile, the total weights *α*_S_ +*β*_S_ and *α*_AA_ + *β*_AA_ are concentration parameters controlling the amount of information contributed by the priors.

For each codon site *i* = 1, … , *I*, define the collection of 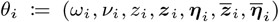 to be the *selection profile* at site *i*, where 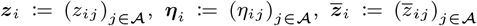 and 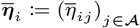. The *fitness profile* **Ψ**_*i*_ is a deterministic function of *θ*_*i*_, with any sample of *θ*_*i*_ inducing a corresponding sample of **Ψ**_*i*_. Inference for all parameters can be performed using MCMC methods as described below.

## 4 Markov chain Monte Carlo

The selection profile *θ*_*i*_ defined in Section 3.2 contains both the underlying model parameters and their deterministic transformations. For posterior inference, it suffices to sample the underlying parameters 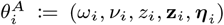, from which the remaining quantities can be obtained directly from (9). (We use the superscript *A* to distinguish this *ancillary parameterization* from the alternative *sufficient parameterization* introduced in 4.2.) Letting 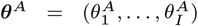, Bayesian inference of selection parameters is performed by obtaining the joint posterior distribution *P* (***θ***^A^ | *Y, g*) of the selection profiles ***θ***^A^ := (*θ*^A^, … , *θ*^A^) given the observed sequences *Y* and the tree *g*. Under the model specified by (1) and (2), pruning can be used to evaluate the likelihood with respect to ***θ***^A^, and standard MCMC procedures applied to sample from the posterior distribution.

### 4.1 Gibbs Sampler

For the model defined by (5)-(9) above, it is natural to perform MCMC by constructing a Gibbs sampler (Brooks et al. 2011) that updates the parameters and latent indicators conditionally. For a given site *i*, the updating of the inclusion indicators and coefficients alternates between two steps, drawing from the conditional distributions:

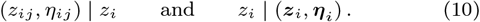

The full description of this Gibbs sampler is given in Appendix A.

Unfortunately, the straightforward Gibbs sampler described by (10) suffers from slow mixing. One cause of this can be seen by considering transitions out of the set {***θ***^A^ : *z*_*i*_ = 0}. When *z*_*i*_ = 0, the parameters *z*_*ij*_ ‘s and *η*_*ij*_ ‘s no longer appear in the likelihood (and therefore the acceptance probability). Consequently, when *z*_*i*_ = 0, the sampler updates these parameters by drawing from their priors, and they may drift away from the support of the data. When a subsequent proposal of *z*_*i*_ = 1 then attempts to reactivate the *z*_*ij*_ ‘s and *η*_*ij*_ ‘s, these values rarely fit the data well. As a result, such proposals tend to be rejected with high probability and the chain becomes effectively trapped at *z*_*i*_ = 0. This behavior is shown in Figure 1. We have found this slow mixing to be a significant practical barrier to analyzing realistic sequence lengths and tree sizes. In the following section, we provide an acceleration strategy that significantly mitigates this problem.

**Figure 1:**
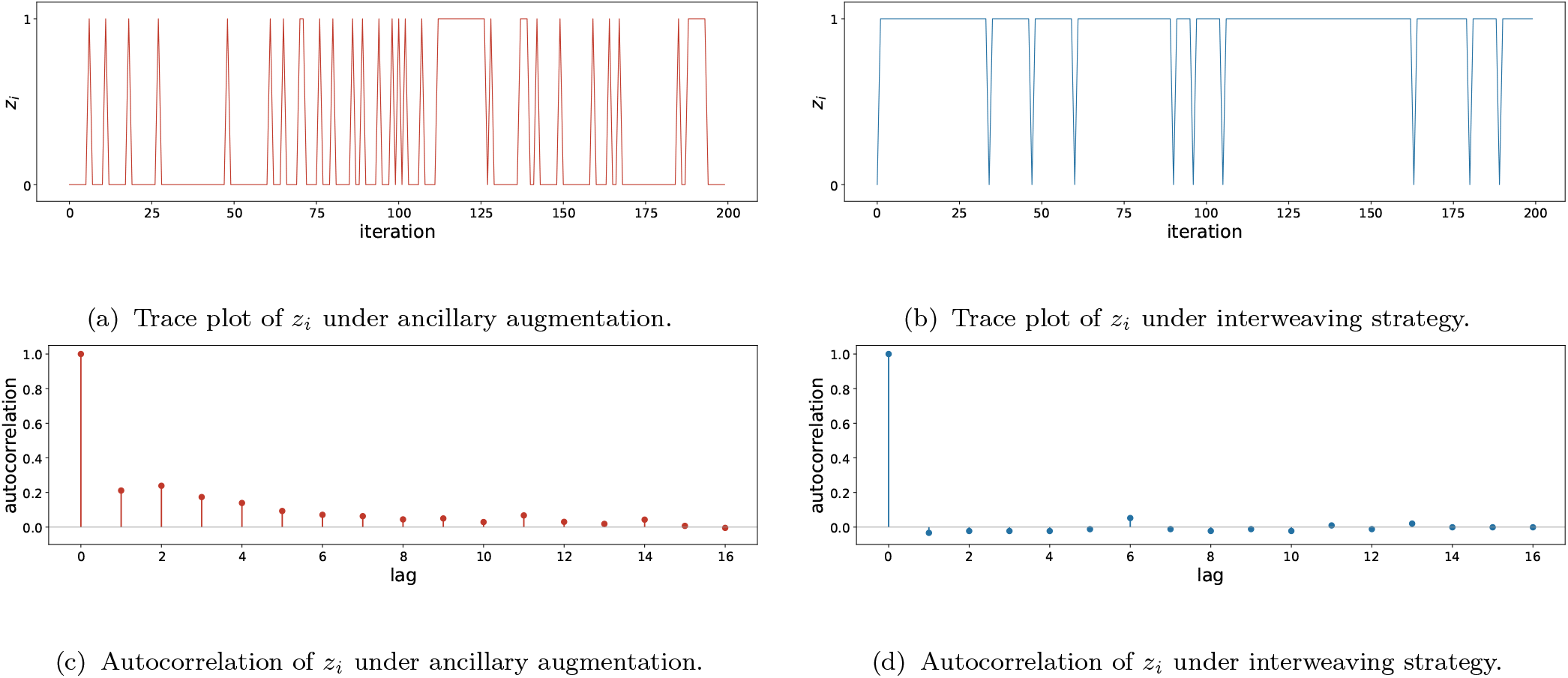
Mixing at a site under selection. Mixing behavior of the site inclusion indicator under the straightforward Gibbs sampler (ancillary parameterization), compared with the behavior under the interweaving strategy, for a single site simulated along an 80-leaf tree with 10 of 20 amino acids truly under selection. Since 10 amino acids are under selection, the posterior inclusion probability *π*_*i*_ := *P* (*z*_*i*_ = 1 | *Y, g*) for this site is high, so a well-mixing chain should occupy *z*_*i*_ = 1 in most iterations. (a) Under the original Gibbs sampler (AA), the chain instead spends most iterations trapped at *z*_*i*_ = 0, with an acceptance rate for the transition to *z*_*i*_ = 1 of only 0.26 (mean recurrence time for state 1 of 4.84 iterations). The apparent frequent switching in (a) therefore reflects repeated trapping in the low-posterior state rather than good mixing. (b) Under the interweaving strategy, the chain correctly remains at *z*_*i*_ = 1 for most iterations; its occasional excursions to *z*_*i*_ = 0 last a single iteration (acceptance rate 100% for 0 → 1 transitions, mean recurrence time for state 1 of 1.00). Correspondingly, the autocorrelation of *z*_*i*_ drops to near zero after lag 1 under ASIS (d), compared with the persistent autocorrelation under AA (c).

### 4.2 Reparameterization

One way to understand – and address – the slow mixing problem is to recognize that the inclusion indicator variables and the selection coefficients in the model (5-9) can be *reparameterized* into a slightly different but equivalent form. In particular, the parameterization (5-9) can be viewed as an *ancillary augmentation* (Yu and Meng 2011; Li and Schmidler 2026) (Figure 2a), suggesting an alternative parameterization, the *sufficient* augmentation. That is, the hierarchical mutationselection model (5-9) can be equivalently rewritten as

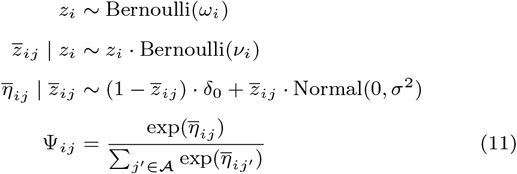

**Figure 2:**
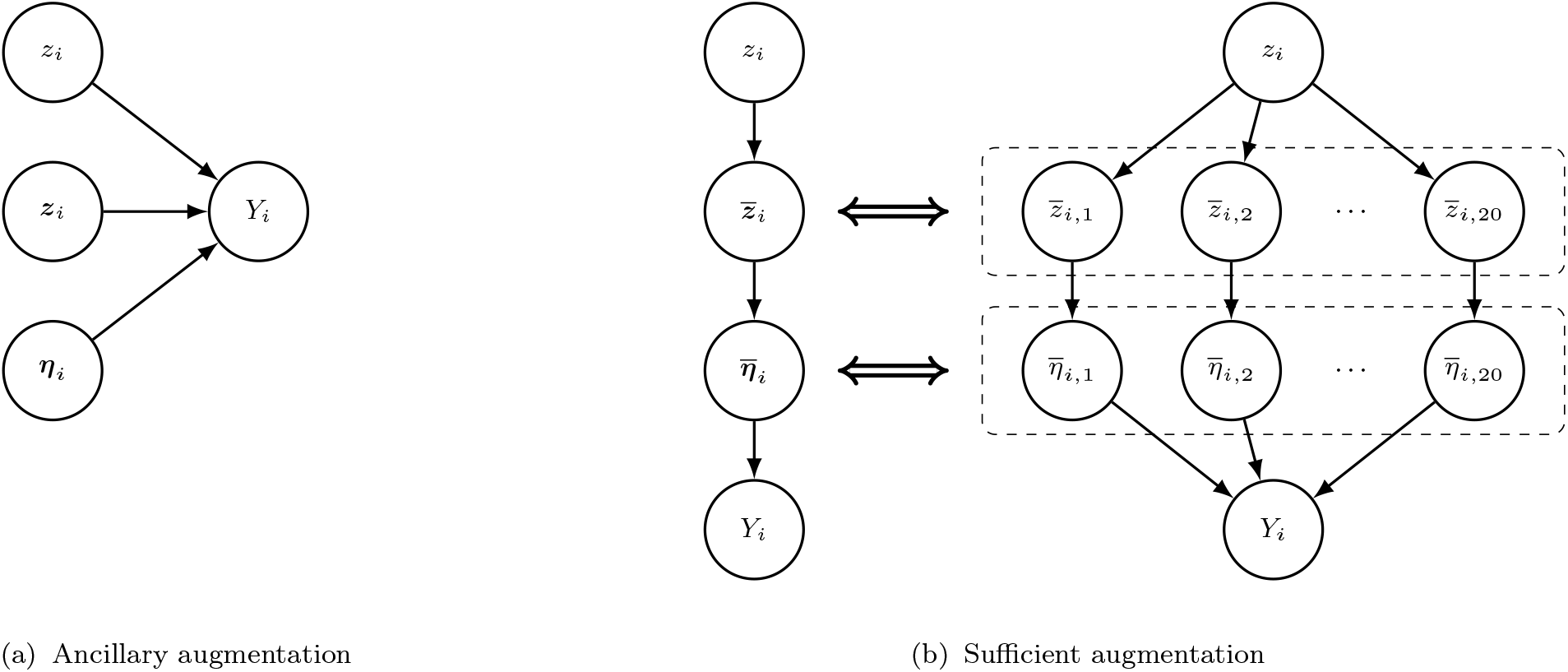
A graphical representation of the BVS model under (a) ancillary augmentation and (b) sufficient augmentation.

The parameterization (11) is called the sufficient augmentation (Yu and Meng 2011; Li and Schmidler 2026) because the augmented parameters 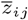 and 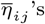 are sufficient statistics for *z*_*i*_ (Figure 2b). Under this reparameterization, deriving the full conditionals yields a Gibbs sampler different from the one specified by (10), which iteratively draws from the conditional distributions:

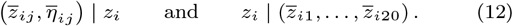

Unfortunately, the sufficient augmentation (11) also exhibits a different, complementary slow mixing transition. In this case, the site inclusion indicator *z*_*i*_ cannot be deactivated (transition from *z*_*i*_ = 1 to *z*_*i*_ = 0) when any *z*_*ij*_ = 1. Therefore, the chain becomes trapped at the state {***θ*** : *z*_*i*_ = 1} until all lower level inclusion variables are simultaneously equal to 0 (*z*_*ij*_ = 0 for all *j*), which is rare since each *z*_*ij*_ has an independent – if small – probability of inclusion. The resulting slow transition leads to high autocorrelation and slow mixing, as shown in Figure 3.

**Figure 3:**
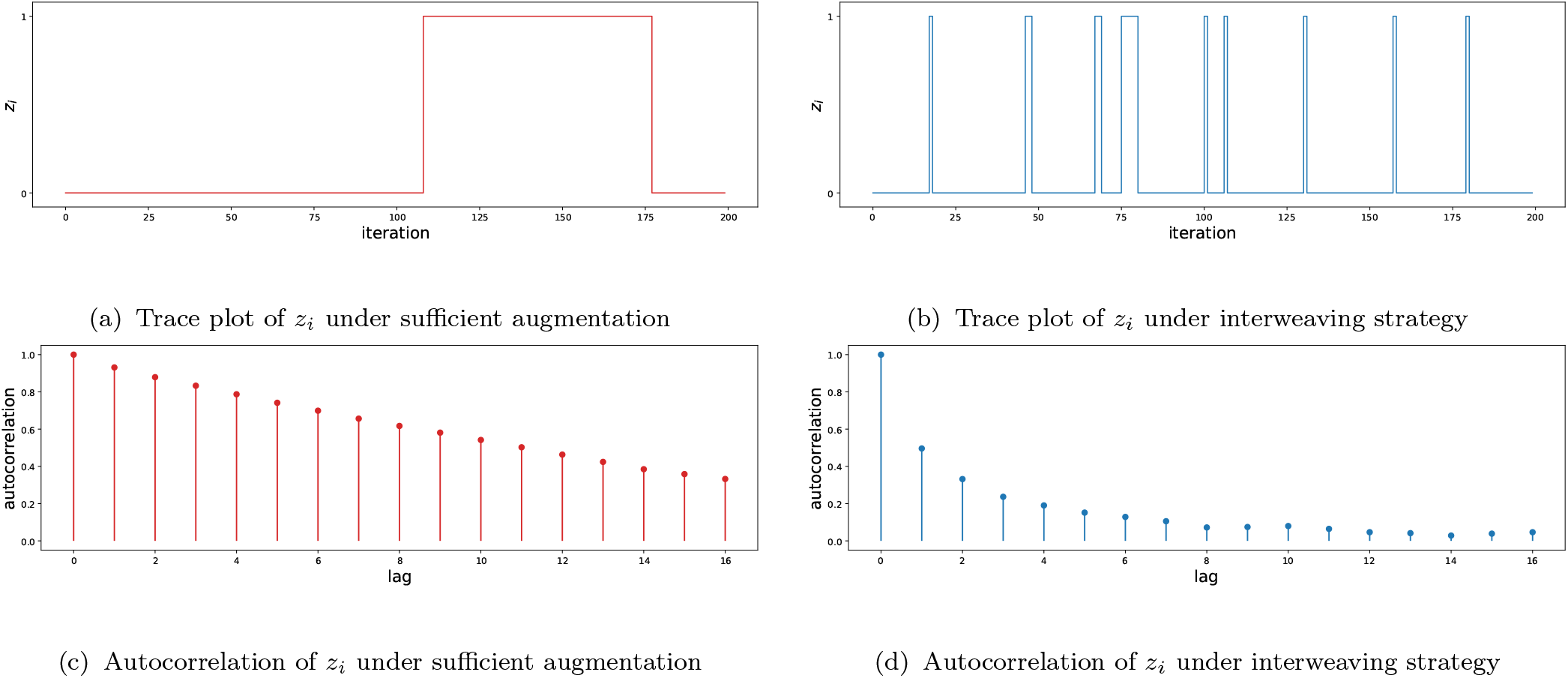
Mixing at a neutral site. Mixing behavior of the site inclusion indicator under the sufficient augmentation compared with the interweaving strategy, for a single fully neutral site (with no amino acids under selection) simulated along an 80-leaf tree. Under the sufficient augmentation, the acceptance rate for 1 → 0 transitions is 0.07, and the mean recurrence time for state 0 is 276.24. Under the interweaving strategy, the transition acceptance rate is 0.47, and the mean recurrence time for state 0 is 30.72.

### 4.3 An accelerated Gibbs sampler

The hierarchical inclusion structure (*z*_*i*_, *z*_*ij*_) of our model is an instance of the general hierarchical group variable selection problem studied in (Li and Schmidler 2026), which shows that component-wise Gibbs/Metropolis samplers built from either parameterization alone often suffer slow mixing due to one of the low-probability transitions described above. Li and Schmidler (2026) resolve these issues by formulating the problem as an instance of the *ancillarity–sufficiency interweaving strategy* (ASIS) (Yu and Meng 2011), which interweaves updates under both parameterizations within a single iteration, and shows that this regularly outperforms either strategy individually as well as the combined strategy that alternates between the two (see (Yu and Meng 2011; Li and Schmidler 2026) for details). We adopt this interweaving strategy here.

Following the general construction of (Li and Schmidler 2026), a single iteration for site *i* proceeds by sampling

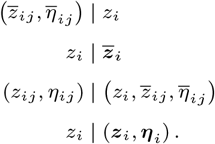

The first step updates the lower-level indicators and coefficients jointly conditional on *z*_*i*_ via a Metropolis step, while the second step resamples the group indicator conditional on the updated lowlevel indicators and coefficients. Note that *z*_*i*_ = 1 with probability one whenever ∑_*j*_ *z*_*ij*_ *>* 0. Otherwise, *z*_*i*_ is drawn from the conditional Bernoulli distribution:

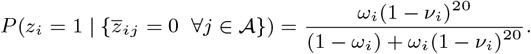

The third step carries these values into the ancillary parameterization: if *z*_*i*_ = 1, we set 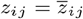 and 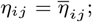 while if *z*_*i*_ = 0, we draw *z*_*ij*_ ∼ Bernoulli(*v*_*i*_) and *η*_*ij*_ ∼ N(0, *σ*^2^) directly from their prior distributions since *z*_*ij*_ and *η*_*ij*_ are independent of *z*_*i*_. The final step updates the group indicator again under the ancillary parameterization, by proposing 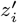 from its prior and accepting with probability min(1, *ρ*), where *ρ* is a ratio of marginal likelihoods under the rate matrices resulting from 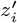 versus *z*_*i*_. The marginal likelihoods can be calculated directly using pruning.

Figure 1(b) and 1(d), 3(b) and 3(d), and 4 show the effects of applying this interweaving strategy. Figure 4 shows the convergence of the running estimate 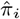 of the posterior probability of site *i* being selected, at a neutral site and at a site with several amino acids under selection. In each case, ASIS outperforms the SA and AA parameterizations, converging within a few hundred iterations at the neutral site and a few thousand iterations at the selected site, whereas the SA and AA schemes require many more iterations to do so. We see a significant improvement in convergence of the time averages (Figure 4) as well as in the frequency of transitions in the traceplots (Figures 1(a,b) and 3(a,b)) and estimated autocorrelation (Figures 1(c,d) and 3(c,d)). In practice, we have found that without this acceleration it is very difficult to obtain convergence of selection estimates on full-length sequences (Table 1).

**Table 1.** Effective sample size per unit CPU time of the resulting MCMC algorithm using different parameterizations for the 3 alignments analyzed in Section 5.

|  | SA | AA | ASIS |
| --- | --- | --- | --- |
| GLOBIN17-144 | 8.4 | 14.4 | 174.9 |
| SDHB26-168 | 5.9 | 11.1 | 131.0 |
| BLAC50-260 | 4.1 | 1.8 | 52.6 |

**Figure 4:**
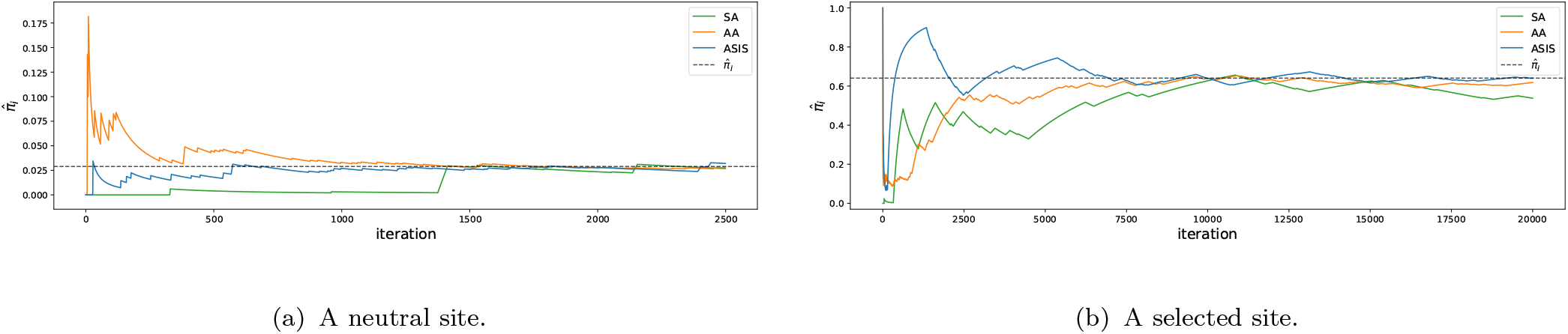
Running estimate (under SA, AA, and ASIS) of the posterior probability *π*_*i*_ of a site being under selection, shown for (a) a neutral site; and (b) a site where four of the twenty possible amino acids are under selection. Dashed lines show the true probability under convergence, verified by multiple-chain convergence diagnostics and averaging. The ASIS estimate converges fastest in both settings, with SA and AA converging substantially slower, particularly under selection.

## 5 Results

To evaluate the impact of the BVS approach on selection detection, we first consider simulated datasets where the ground truth can be known exactly. We examine the operating characteristics of detection performance measure, being careful to account for the impact of selection signal strength. We then explore the impact on identifying selection in real sequences by re-analyzing three datasets from the literature: the GLOBIN17-144 alignment (17 vertebrate sequences of *β*-globin gene, length 144 codons) used by (Yang et al. 2000; Rodrigue et al. 2010), SDHB26-168 (26 sequences of succinate dehydrogenase subunit B, 168 codons) used by (Kullberg et al. 2006; Rodrigue et al. 2010), and BLAC50-260 (50 class A *β*-lactamase sequences, 260 codons) from the alignment distributed with phydms (Hilton et al. 2017) together with the deep mutational scan of TEM-1 *β*-lactamase reported by (Stiffler et al. 2015).

### 5.1 Simulation studies

To evaluate the accuracy of our approach when ground truth is known, we simulated multiple data sets, each representing a multiple sequence alignment input to a phylogenetic analysis. For each dataset, we simulated the full evolutionary history as follows: first, a phylogenetic tree was generated by recursively selecting a leaf uniformly at random to bifurcate until the desired number of taxa was reached, and branch lengths were drawn independently from an exponential distribution. Next, selection profiles for each codon site were sampled independently from the prior distribution with *α*_S_ = 35, *β*_S_ = 65, *α*_AA_ = 15, *β*_AA_ = 85, *σ* = 10. Finally, an ancestral sequence of length 168 codons was drawn from the stationary distribution (3) and the full evolutionary history of sequences along the tree was simulated according to the mutationselection model (2). To explore the effects of the size of the dataset, datasets were generated with leaf counts ranging from 8 to 64.

Since a primary goal of mutation-selection modeling is to identify the presence of selection, we wish to assess the operating characteristics of detection performance measures (sensitivity, specificity, precision) as the underlying selection signal strength varies. Some care is required in doing so, as multiple variables combine to determine the amount of information available for detection in the dataset: the number of sequences observed, the branch lengths and mutation rates, and the magnitude of the selection effects. For example, while sensitivity might be expected to scale with the number of sequences observed, the leaf count alone is a poor predictor if branch length is not controlled for (Figure 5). Similarly, (mean) branch-length is inadequate without accounting for the number of leaf sequences. Total branch length captures both, but fails to account for differences in mutation rates. Expected number of substitutions along the tree (per codon site) captures the joint effects of total branch length and mutation rate, and therefore seems to be the most direct and invariant measure of information content; we find that datasets with comparable mutations per codon site achieve comparable sensitivity in detecting positive selection regardless of leaf count, as shown in Figure 5. However, mutation count is also confounded with selection strength, with stronger selection often inhibiting mutation due to fixation, even while stronger selection signals are expected to be easier to detect. Thus to assess the operating characteristics of the BVS model, we further stratify simulated sites into weak, moderate, and strong selection regimes defined by realized fitness contrast Δ**Ψ**_*i*_ := max_*j*_ Ψ_*ij*_ − min_*j*_ Ψ_*ij*_ values of (0, 0.4], (0.4 − 0.75], and *>* 0.75 respectively. Figure 6 shows the distribution of total variation (TV) distance between the true profile and the uniform profile obtained for each selection regime. Separately we also evaluated specificity on truly neutral sites (*z*_*i*_ = 0).

**Figure 5:**
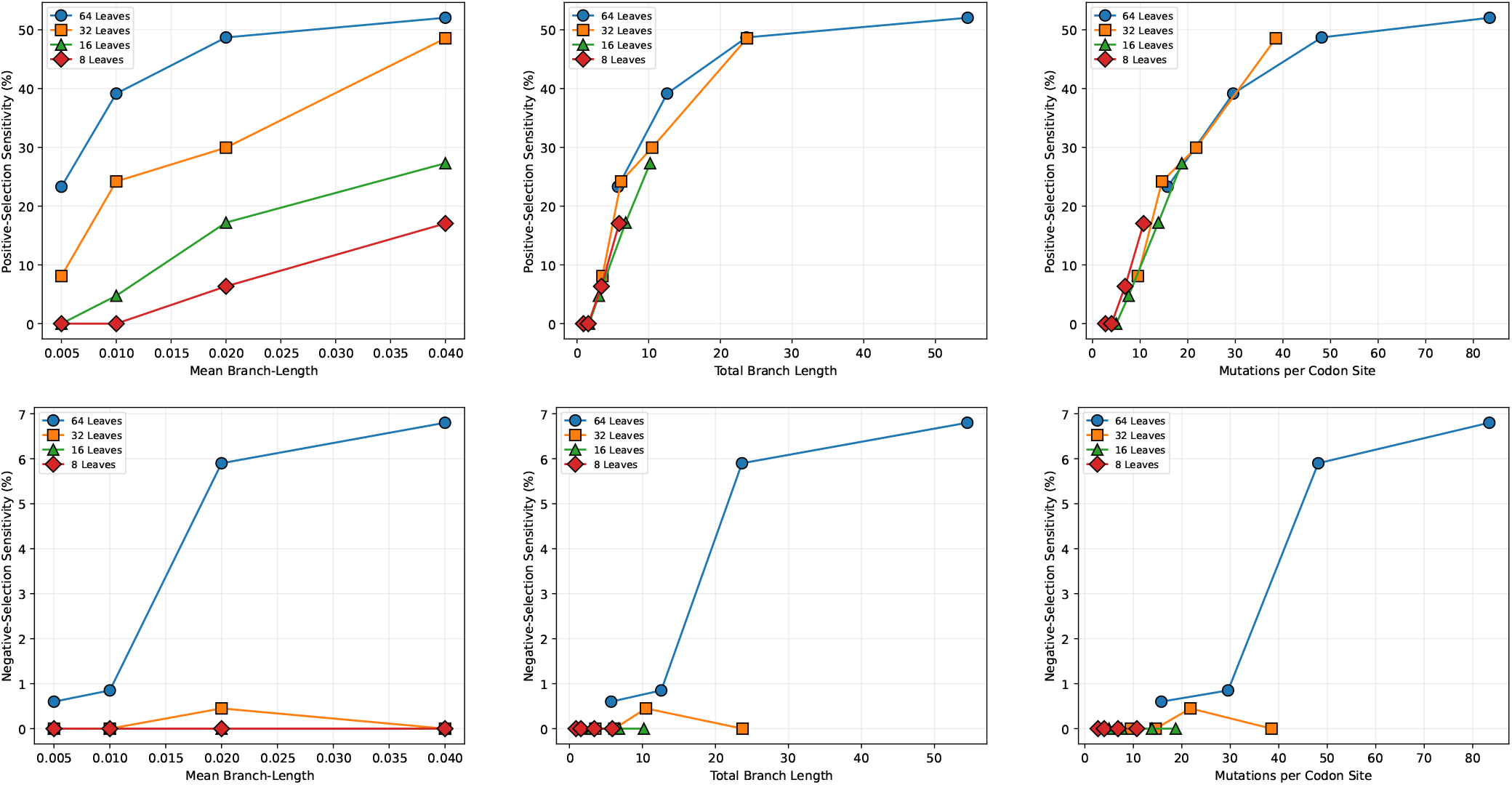
Mutations per codon site determines detection sensitivity better than leaf count or branch-length rate alone. Shown are sensitivities for detecting positive (top) and negative (bottom) amino-acid-specific selection, as a function of mean branch-length (left), total branch length (middle), and mutations per (codon) site (right). Different lines/colors indicate leaf count (8–64). Points represent mean of two replicate datasets at each leaf count+branch-length setting. Sensitivity for sites under negative selection remains near zero except at higher leaf counts and branch-lengths/mutation counts (bottom row), reflecting the difficulty of inferring at high confidence from the absence of observed mutation events.

**Figure 6:**
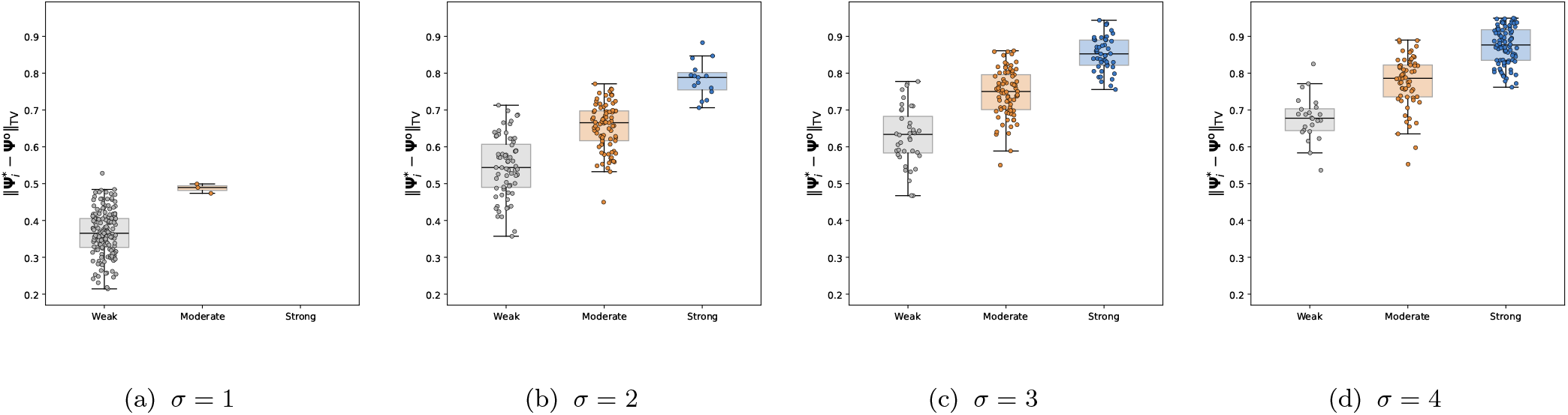
Distribution of the total variation distance between each site’s true (simulated) profile 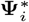 and the uniform profile **Ψ**^o^ obtained under each selection regime.

Figure 7 shows the true positive rates (sensitivity) and false positive rates (1-specificity) on the simulated datasets as a function of the posterior inclusion probability threshold, with curves for independent datasets colored by mean mutations per codon site. We see that sensitivity increases with increasing selection strength, as well as monotonically with mutation count within each selection strength regime. Sites under strong selection are detected with near-perfect sensitivity at moderate mutation counts, while sites under weak selection require substantially more mutations to be reliably detected. Even sites under strong selection may be difficult to detect at low expected mutation count, for example when multiple nucleotide changes are required to reach the selected amino acid from the ancestral sequence codon, or the elapsed evolutionary time (branch length) is too short. In contrast, the false positive rate at neutral sites (Figure 7d) falls sharply once the threshold exceeds the prior inclusion probability, and remains low independently of the amount of information. The ability to continuously tune sensitivity and specificity by choosing the prior inclusion probability threshold is a direct consequence of the BVS framework, which estimates the inclusion probability separately from the selection coefficient.

**Figure 7:**
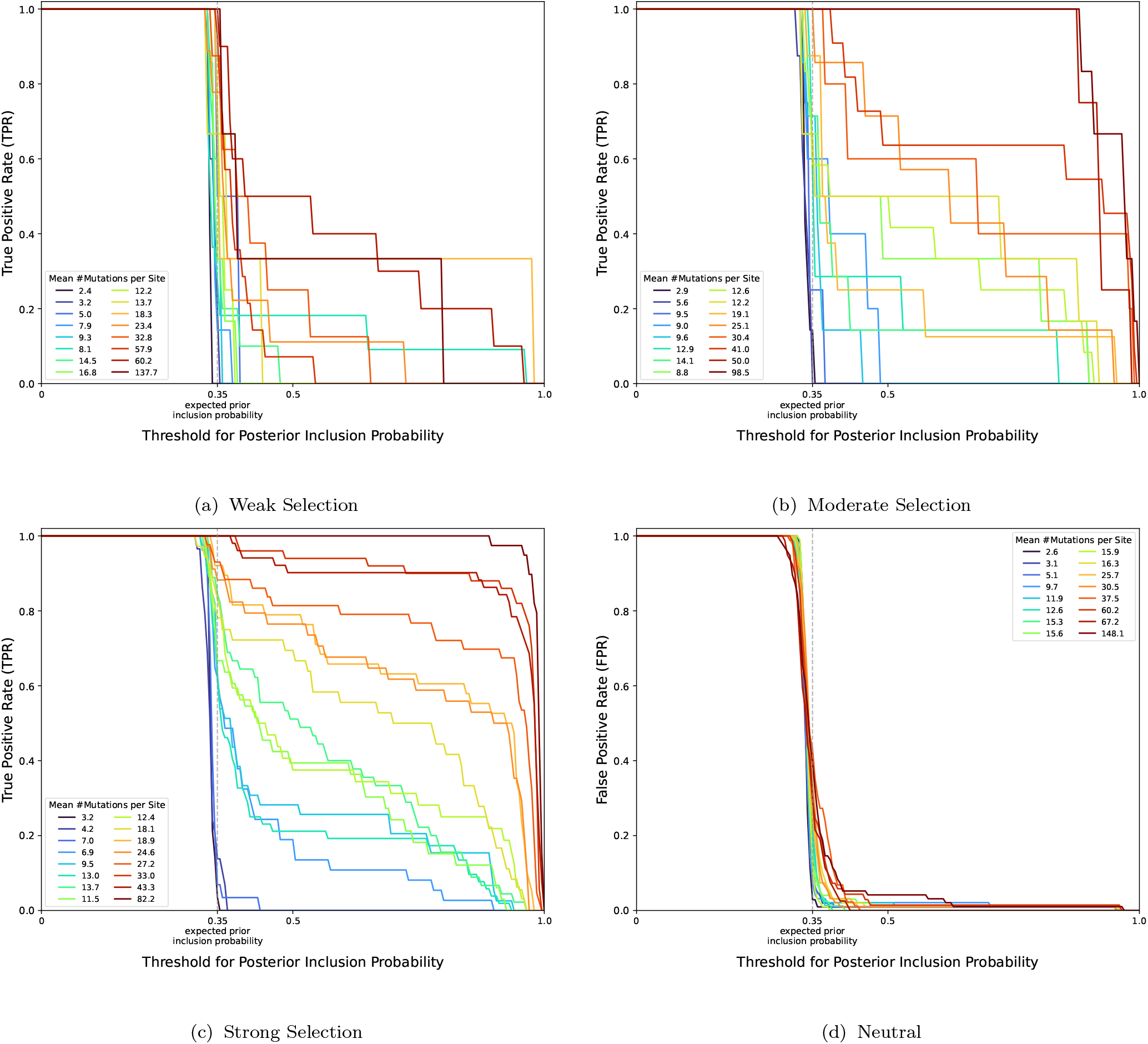
Site-level sensitivity and specificity of the BVS model as a function of the posterior inclusion probability threshold.

#### Comparing selection vs. shrinkage

We next compared two strategies for controlling the complexity of the site-heterogeneous mutation-selection model: the *model selection* approach described in Section 3.2, and the *shrinkage estimation* approach using DP priors, as discussed in Section 2.3. Under model selection, complexity is controlled by explicitly excluding parameters from the model, while under shrinkage estimation, complexity is instead controlled by pooling information across sites. These two strategies rely on different structural assumptions about the data; here we explore which may be preferable under different selection-strength regimes. To this end, we simulated multiple phylogenetic datasets with trees generated as described above, but with selection profiles for each codon site sampled under the *dense model* (Halpern and Bruno 1998) at four signal strengths *σ* ∈ {1, 2, 3, 4} before simulating the evolution of sequences along the tree. For a range of choices of (*α*_S_, *β*_S_) and (*α*_AA_, *β*_AA_) spanning a corresponding range of prior expectations (E [*ω*] , E [*v*]) =: (*p*_S_, *p*_AA_) ∈ (0, 1)^2^, we fit the mutation-selection model to the sampled dataset using the BVS approach, and computed the L1 distance between the posterior mean profile and the ground truth. Figure 8 shows this divergence averaged across sites and smoothed over the (0, 1)^2^ grid. The divergence is also computed for the posterior obtained under the DP-mixture (Lartillot and Philippe 2004; Rodrigue et al. 2010; Lartillot et al. 2013; Rodrigue and Lartillot 2014) prior using the following parameters (identical to (Lartillot and Philippe 2004)): *c* ∼ Uniform [0.001, 1.0], *G*_0_ = Dirichlet (1, … , 1), **Ψ** ∼ DP (*c, G*_0_).

**Figure 8:**
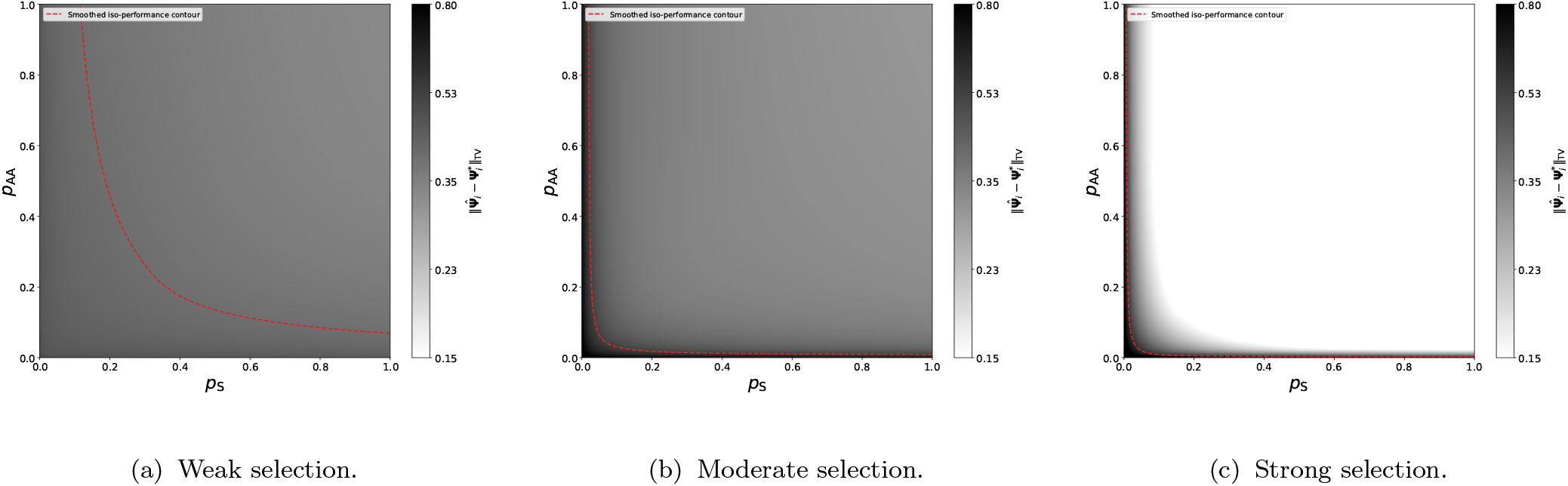
Total variation distance between estimated and true site-specific amino acid fitness profiles under the BVS prior as a function of the site-level prior inclusion probability *p*_S_ and the amino-acid-level prior inclusion probability *p*_AA_ (with *α*_*S*_ + *β*_*S*_ and *α*_*AA*_ + *β*_*AA*_ held fixed) for datasets obtained under selection coefficients simulated from independent normal distributions with variance *σ* = 1, 2, 3, 4. The red dashed curve is the iso-performance contour indicating the (*p*_S_, *p*_AA_) values at which BVS matches the performance of the DP mixture prior, obtained by smoothing MCMC evaluations across the grid. Below and left of the contour, the DP-mixture prior outperforms BVS; above and right, BVS outperforms the DP-mixture prior. In the presence of weak selection, the BVS approach is largely insensitive to (*p*_S_, *p*_AA_), while under strong selection BVS outperforms the DP-mixture unless the prior inclusion probabilities are set to very small values. In the moderate selection regime where the signal in the data is less clear, the differences are especially pronounced, with the divergence achieved by the DP-mixture prior indicating over-shrinkage toward shared cluster profiles.

Figure 8 shows the contour of (*p*_S_, *p*_AA_) at which BVS achieves performance identical to that achieved by the DP-mixture prior. Note that the dense mutation-selection model is recovered as a limiting case of BVS as the site- and amino-acid-level prior inclusion probabilities *p*_S_, *p*_AA_ approach 1 (equivalently, as the hyperparameters *β*_S_, *β*_AA_ → 0). At these hyperparameter values, we therefore expect BVS to outperform the DP-mixture approach on data simulated from the fully dense model when data is sufficiently informative, which is indeed observed in Figure 8. The amount of information required for this to occur, however, depends on the magnitude of the underlying selection effect. For weak selection, the true fitness profiles lie closer to the uniform distribution (Figure 6), and the distinction between a neutral site and one under selection is difficult to resolve without large amounts of data (mean mutations/site). In the absence of sufficient signal, the BVS prior conservatively identifies sites as neutral rather than inferring selection that is not well supported by the observed data at that site. The DP-mixture prior, in contrast, pools information from other sites assigned to the same mixture components. While this could improve estimation when several sites genuinely share the same selection profile, such as sites selecting for general sidechain properties such as hydrophobicity or volume, it can also introduce errors when such cross-site similarity is not present (see Figure 9a).

**Figure 9:**
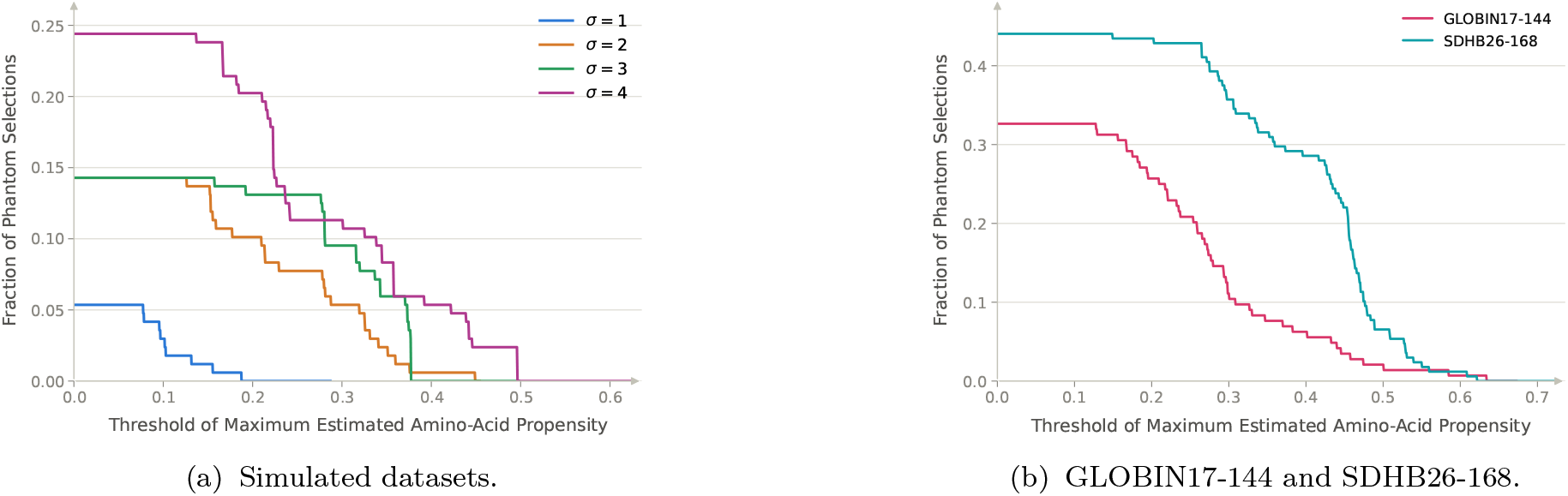
Phantom selections under the DP-mixture prior. Fraction of sites whose DP-mixture estimate is a phantom selection (highestpropensity amino acid absent from the observed sequences) with 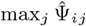 exceeding the threshold *t*, obtained using the DP-mixture prior approach on: (a) data simulated from the dense model with *σ* = 1, 2, 3, 4, and (b) GLOBIN17-144 alignment and SDHB26-168 alignment. Both GLOBIN17-144 and SDHB26-168 exhibit substantially higher fraction of phantom selection under the DP-mixture prior than those observed in the simulated datasets. The BVS approach never identifies any phantom selections in any of the analyzed datasets.

*Phantom selection:* In fact, the DP-mixture prior exhibits a highly undesirable property: with some frequency it identifies site-amino acid pairs as being under positive selection, *even when that amino acid has never been observed to occur at that site*. We refer to a site whose posterior-mean highest-propensity amino acid is never observed among its own tip sequences as a *phantom selection*. Upon inspection we find that, unsurprisingly, these phantom selections are closely related to strong selection at *other* sites. This is a direct consequence of the DP-mixture pooling information across sites: when a cluster contains one or more sites under strong selection for a particular amino acid, that amino acid’s fitness is driven strongly upward for the whole cluster, and any other site sharing the cluster inherits this preference in its own selection estimates, even if that amino acid never occurs at that site in the observed sequences. This demonstrates both a strength and a weakness of sharing information across sites. While this can be beneficial when identifying general fitness based on broad structural compatibility or hydrophobicity/philicity, when fitness profiles are specific to the local environment or functional role of a given site, such sharing can be counterproductive, resulting in the introduction of errors (phantom selections). Sites with low signal – either due to weak selection or limited data – are especially susceptible, since the likelihood contributes little to resist being pulled toward the cluster’s dominant preference. The result is an artifact of information borrowing rather than a signal actually present in the data at the affected site. This suggests that the DP-mixture formulation is less suitable for identifying truly site-specific selection effects.

To quantify how often these phantom selections occur and how strongly the DP-mixture estimates them to be under selection, we define the estimated propensity vector 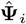 to be a *phantom selection* at threshold *t* if the amino acid having the highest propensity is not observed among the leaf sequences at site *i* and 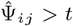. Figure 9a shows the resulting fraction of phantom selections as a function of the threshold *t* on the simulated datasets. The BVS approach never identifies a single phantom selection on any of the datasets.

### 5.2 GLOBIN17-144 dataset

Figure 10 shows the result of applying the BVS model to the GLOBIN17-144 alignment. We see that BVS recovers a small, sparse set of high posterior inclusion probability site+AA combinations: 13W, 16V, 26L, 30L, all with 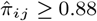, where 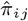 denotes the MCMC estimate of 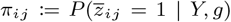. These findings are consistent with the strong, unambiguous selection for these residues in the fit by the dense model (using a DP-mixture prior) of (Rodrigue et al. 2010). A second group of site+AA pairs (6K, 15K) is assigned moderate inclusion probability 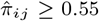, reflecting comparatively weaker evidence of selection at these sites. The majority of the remaining sites exhibit posterior inclusion probability below 0.5 despite occasionally large point estimates of the actual selection coefficient *η*_*ij*_ (e.g., 5E, 22G, 25A, 28R). At these sites, the low posterior inclusion probability (i.e. 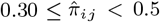, close to or slightly above the prior mean 0.35) leads to the sites being excluded by the BVS prior to preserve sparsity. This demonstrates the advantage of the BVS framework discussed in Section 3.1: the estimates of inclusion probability and selection coefficient magnitude at a given site need not agree, and their divergence can be informative. We also observe phantom selections in this dataset when using DP-mixture priors (Figure 9b). In particular, at 41 of the 144 sites the amino acid with the highest estimated propensity, 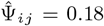, does not appear in any of the observed sequences at that site.

**Figure 10:**
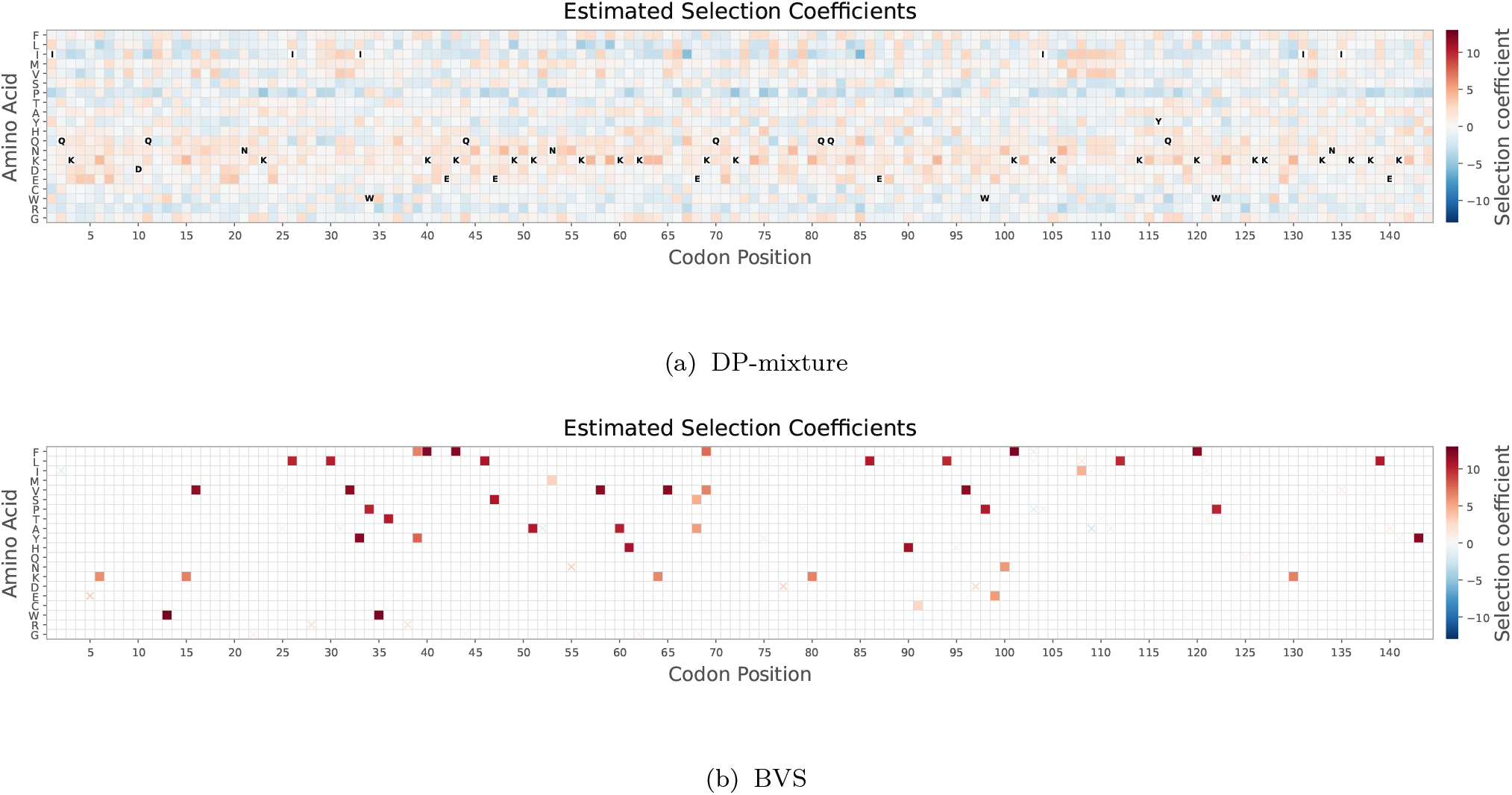
The posterior mean selection coefficients *η*_*ij*_ for the GLOBIN17-144 dataset. Color encodes the sign and magnitude of the estimated selection coefficient 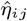 for amino acid *j* at codon site *i* (red: favored; blue: disfavored). (a) Using DP-mixture priors. For phantom selections, the square for the highest-propensity amino acid is left uncolored and marked with its amino-acid letter. 47 of the 144 sites exhibit phantom selections. (b) Using BVS priors. Markers are rendered as filled squares when 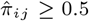 and as crosses (*×*) otherwise, with color encoding the sign and magnitude of 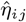.

### 5.3 SDHB26-168

Application of the BVS model to the SDHB26-168 alignment identifies a sparse set of residues with large estimated selection coefficients, several of which coincide with sites independently implicated in SDHB structure and function (Table 2). Most notably, 7 of the 63 residues identified by the BVS analysis (C93, C98, C101, C113, C186, C189, and C196) are conserved cysteines that directly coordinate the SDHB iron-sulfur clusters (Vranken et al. 2015; Saxena et al. 2016). Consistent with the importance of these residues, a recent cellular complementation study found loss-of-function phenotypes for the SDHB variants C93F, C101S, C113G, C186S, and C189R (Lee et al. 2026). For C196, previous experimental work showed that the C196Y variant reduces SDHB protein stability through a shortened protein half-life (Yang et al. 2012). The same complementation study also found that A102T retained approximately 20% of wild-type SDH activity, consistent with partial functional impairment (Lee et al. 2026). Finally, the F146L substitution was shown experimentally to greatly diminish SDHB–SDHAF1 binding (Maio et al. 2016). Together, these results provide independent functional support for several of the residues highlighted by the BVS analysis, while illustrating that the inferred selection coefficients recover sites associated with distinct biochemical mechanisms. Under the DP-mixture prior, SDHB26-168 suffers even more from phantom selections (Figure 9b) with 50% of the sites having a highest-propensity amino acid with 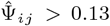 that does not appear in any of the observed sequences at that site.

**Table 2.** Independent biochemical and functional evidence for SDHB residues identified by the BVS analysis. Shown are human SDHB residues with large estimated selection coefficients that coincide with sites independently implicated in SDHB structure or function. Remarks summarize the relevant biochemical or experimental evidence.

| Human SDHB residue | $\hat{\eta}$ | Remark |
| --- | --- | --- |
| C93 | 1.42 | Fe-S ligand; substitutions impair SDHB function |
| C98 | 1.34 | Fe-S ligand; perturbation disrupts Fe-S-dependent SDHB function |
| C101 | 1.25 | Fe-S ligand; C101S produces loss of function |
| A102 | 1.25 | A102T shows partial functional impairment |
| C113 | 1.27 | Fe-S ligand; C113G produces loss of function |
| F146 | 2.25 | F146L impairs SDHB-SDHAF1 binding |
| C186 | 1.35 | Fe-S ligand; C186S produces loss of function |
| C189 | 1.07 | Fe-S ligand; C189R produces loss of function |
| C196 | 1.37 | C196Y reduces SDHB stability (shortened half-life) |

### 5.4 BLAC50-260

This dataset consists of an alignment of 50 *β*-lactamase sequences spanning 260 codons, together with site- and aminoacid-specific preferences derived from deep mutational scanning (DMS) measurements of TEM-1 *β*-lactamase under ampicillin selection (Stiffler et al. 2015; Hilton et al. 2017). DMS couples large libraries of protein variants to functional selection and quantifies changes in variant frequencies by high-throughput sequencing (Fowler and Fields 2014). Following the experimentally informed codon-model formulation of (Bloom 2014; Hilton et al. 2017; Hilton and Bloom 2018), the resulting enrichment ratios are normalized across the 20 amino acids at each site to obtain amino-acid preference vectors on the same probability simplex as the site-specific propensity vectors **Ψ**_*i*_, allowing site-wise comparison between the experimental and phylogenetic estimates. These DMS-derived preferences therefore provide an independent experimental reference for evaluating the inferred site-specific amino-acid preferences. We use the codon alignment and the DMS data (Stiffler et al. 2015) distributed with phydms (Hilton et al. 2017), remove the three alignment columns containing gaps, and estimate a phylogenetic tree and the General Time Reversible (GTR) mutation model parameters from the remaining 260 columns, which are then held fixed in the subsequent mutationselection analyses. Sites are reported in the standard Ambler numbering for class A *β*-lactamases (Ambler et al. 1991).

The BVS and DP-mixture priors yield markedly different estimates of site-specific amino-acid preferences (Figure 11). Across the 260 sites, the median maximum estimated aminoacid weight, 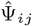, is 0.053 under BVS, close to the uniform value of 0.05, with 23% of sites having a maximum weight within 0.001 of 0.05. In contrast, the corresponding median under the DP-mixture prior is 0.119, with a minimum of 0.081. Thus, BVS estimates approximately uniform amino-acid preferences at many sites where the posterior provides insufficient support for substantial deviation from neutrality, whereas the DP-mixture estimates are more strongly nonuniform across this dataset.

**Figure 11:**
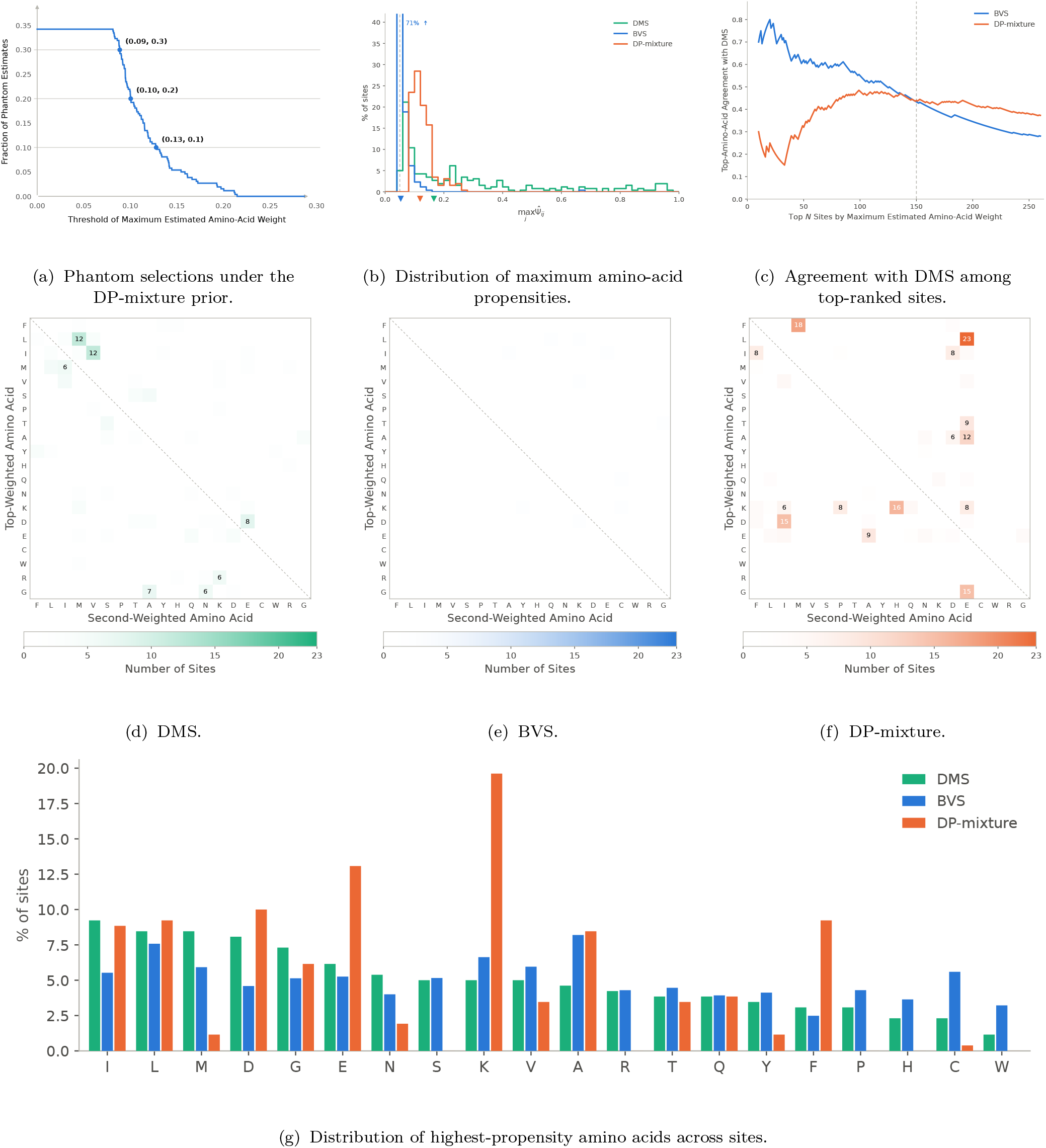
Comparison of BVS and DP-mixture estimates for the BLAC50-260 dataset using amino-acid preferences measured by deep mutational scanning (DMS) as an independent experimental reference. (a) Fraction of phantom selections under the DP-mixture prior as a function of the threshold applied to the maximum estimated amino-acid weight, 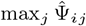. (b) Distribution of 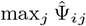 across the 260 sites for BVS and the DP-mixture prior, together with the corresponding maximum DMS preferences. Markers below the horizontal axis indicate medians, and the dashed vertical line denotes the uniform amino-acid weight, 1*/*20 = 0.05. (c) Agreement with DMS as a function of the number *N* of highest-ranked sites. Sites are ranked separately for each model in decreasing order of 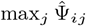, and agreement is the fraction of the top *N* sites for which the model’s highest-propensity amino acid agrees with the DMS-preferred amino acid, arg 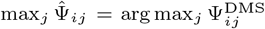. (d–f) Joint distribution of the identities of the top- and second-highest-propensity amino acids across sites according to (d) DMS, (e) BVS, and (f) the DP-mixture prior. Each cell gives the number of sites having the corresponding pair of top- and second-highest-propensity amino acids. (g) Distribution of the identity of the highest-propensity amino acid across sites for DMS, BVS, and the DP-mixture prior. The distribution obtained under the DP-mixture prior shows clear artifacts of over-smoothing by the Dirichlet process, with most probability mass concentrated on several amino acids (K, E, D, L, I, A) and some others (S, R, P, H, W) showing no probability mass at all.

Phantom selections are also observed in this dataset under the DP-mixture prior (Figure 11a). At the 89 sites for which the highest-propensity amino acid is absent from all observed sequences at that site, the corresponding amino acid has a mean DMS preference of 0.059 and a mean DMS rank of 8.7 among the 20 amino acids. In contrast, at the remaining 171 sites, the highest-propensity amino acid has a mean DMS preference of 0.265 and a mean DMS rank of 2.9. Thus, the amino acids identified by phantom selections have substantially lower DMS preferences and less favorable DMS ranks than the highest-propensity amino acids at non-phantom sites. This disagreement with experimental validation is clear evidence of the DP-mixture estimates being problematic.

This distinction is also apparent when sites are ranked by 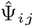 (Figure 11c). Among the top *N* = 20 and *N* = 50 sites, respectively, the model’s highest-propensity amino acid matches the amino acid with the highest DMS preference at 80% and 62% of sites under BVS, compared with 25% and 32% under the DP-mixture prior. The DP-mixture’s agreement exceeds that of BVS only after approximately *N* = 150 sites are retained, by which point the BVS ranking includes many sites with nearly uniform estimated amino-acid preferences.

Examining the identities of the two most strongly preferred amino acids as a pair provides additional perspective (Figure 11d– 11f). The DMS measurements themselves exhibit a nonuniform joint distribution of the top- and second-highest-propensity amino acids, consistent with some sites favoring multiple amino acids with similar physicochemical properties. Notably, several aminoacid pairs occurring frequently in the DMS measurements, including I–V, L–M, D–E, G–A, and R–K, have similar physicochemical properties (Pakula and Sauer 1989; Betts and Russell 2007) such as hydrophobicity and charge. (Such pairs also tend to be relatively exchangeable in empirical protein substitution matrices (Henikoff and Henikoff 1992).) Thus, nonuniformity in the joint distribution of the two highest-preference amino acids is not unexpected on biochemical grounds. However, the DP-mixture estimates are substantially more concentrated on particular amino-acid pairs than the DMS measurements, whereas the BVS estimates exhibit a more diffuse pattern. This greater concentration on particular pairs under the DP-mixture is consistent with its sharing of amino-acid preference profiles across sites through mixture components: when the sequence data at a site are weakly informative, assignment to a mixture component causes its estimated preference profile to resemble those at other sites assigned to that component. Consequently, strong preferences identified at informative sites can influence the estimated profile of weakly informative sites elsewhere in the alignment.

Taken together, these results illustrate the principal statistical distinction between the two priors. When there is insufficient evidence in the data for differential amino-acid preferences at a given site, the BVS prior can assign sites to the neutral submodel. In contrast, the DP-mixture prior places no point mass on the exactly uniform neutral profile, instead controlling complexity by sharing preference profiles across sites through its mixture components. As a result, the DP-mixture prior handles sites with insufficient evidence by shrinking them towards the selection profiles of other sites with stronger evidence. In this dataset, BVS yields estimates closer to the uniform neutral profile at many sites, while the sites receiving its strongest estimated amino-acid preferences show substantially greater agreement with the independent DMS measurements than do the corresponding highest-ranked sites under the DP-mixture prior.

## 6 Conclusion

Formulation of the identification of site-specific amino-acid selection in phylogenetic mutation–selection models as a hierarchical Bayesian variable selection (BVS) problem has multiple advantages. By assigning positive prior probability to exact neutrality, point-mass priors at the site and amino-acid levels allow posterior inference to simultaneously estimate both the presence of selection and its magnitude and direction. The resulting posterior inclusion probabilities provide a direct measure for ranking sites and amino acids by evidence of selection, without requiring arbitrary thresholds on estimated selection coefficients. Moreover, the sensitivity and specificity of selection detection can be controlled explicitly through the posterior inclusion probability threshold. When compared with modern Dirichlet-process (DP) mixture priors, even on datasets simulated under the dense mutation–selection model, BVS achieves comparable or smaller estimation errors unless hyperparameters are chosen to require very high levels of sparsity. In addition, DP-mixture priors were observed to produce *phantom selection* estimates, in which amino acid preferences are inferred for residues never observed in the dataset. This is a direct result of the mechanism by which DP-mixtures control complexity by pooling information across sites. Such phantom selection estimates were observed in both simulated and real-world datasets using the DP mixture priors, whereas BVS never produced such artifacts. Although the absence of an amino acid from observed sequences does not guarantee the absence of positive selection, the frequent occurrence of phantom selections under the DP-mixture prior raises significant concerns about whether inferred preferences are supported by site-specific evidence.

One downside of the BVS formulation is the potential increase in computational difficulty arising from the hierarchical inclusion structure. Component-wise Gibbs samplers under the ancillary and sufficient augmentations exhibit slow mixing: the ancillary parameterization can become trapped at state *z*_*i*_ = 0, whereas the sufficient parameterization can become trapped at either state. By interweaving both parameterizations, the ancillarity–sufficiency interweaving strategy (ASIS) substantially improves mixing of the resulting MCMC algorithm, making posterior inference practical for real-world sequence alignments.

Potential limitations of the present approach suggest opportunities for further improvement. For example, the present BVS framework treats amino-acid selection indicators independently within each site, without explicitly accounting for physico-chemical similarities among amino acids. A natural extension would introduce additional levels of the inclusion hierarchy that allow for selection of groups of amino acids with similar properties such as hydrophobicity, charge, or side-chain volume. Such a hierarchical structure could incorporate additional biophysical and biochemical information while preserving the ability to identify selection at both the group and individual amino-acid levels. The BVS framework can also be applied in contexts where the assumptions of the basic mutation-selection models described here are violated, for example in the presence of site-dependent evolutionary processes (see (Li et al. 2026)).

In summary, hierarchical BVS provides an interpretable alternative to shrinkage-based mutation–selection models when the primary objective is to identify which specific sites and amino acids exhibit evidence of selection. By explicitly distinguishing posterior support for selection from its estimated magnitude, BVS avoids assigning unwarranted confidence to large but weakly supported selection coefficients. Reporting posterior inclusion probabilities alongside selection-coefficient estimates allows the practitioner to assess both the strength of an inferred effect and the evidence for its presence.

## AcknowledgmentS

This work was partially supported by NSF grant DMS-2347859.

## A Gibbs Samplers

Throughout, fix a codon site *i* and let A denote the set of 20 amino acids. We write *L*(*Y*_*i*_ | *θ*_*i*_) for the pruning-algorithm marginal likelihood at site *i* under selection profile *θ*_*i*_ (equivalently, under the fitness profile Ψ_*i*_ implied by *θ*_*i*_), and *ω*_*i*_, *v*_*i*_ for the site- and amino-acid-level prior inclusion probabilities at site *i*, as defined in (5-9).

### A.1 Gibbs Update for Inclusion Probabilities

The site- and amino-acid-level inclusion probabilities *ω*_*i*_ and *v*_*i*_ enter the model only through the priors on the inclusion indicators, (5)-(6) and (8). They do not appear in the likelihood *L*(*Y*_*i*_ | *θ*_*i*_). Because the Beta hyperpriors are conjugate to the Bernoulli distributions, both parameters can be updated by exact Gibbs draws that need no likelihood evaluation.

Given the site indicator *z*_*i*_, the full conditional of *ω*_*i*_ is

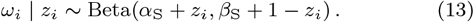

Similarly, given the amino-acid indicators **z**_*i*_ = (*z*_*i*1_, … , *z*_*i*,20_), the full conditional of *v*_*i*_ is

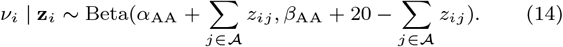

### A.2 Gibbs Update for Fitness Profiles under Ancillary Augmentation

Recall that under the ancillary augmentation, the auxiliary parameters (*z*_*ij*_, *η*_*ij*_)_*j∈A*_ are defined independently of *z*_*i*_ in the prior, and enter the likelihood, through Ψ_*i*_, only when *z*_*i*_ = 1. A single sweep at site *i* updates (*z*_*ij*_, *η*_*ij*_)_*j∈A*_ | *z*_*i*_, and then *z*_*i*_ given the result.

When *z*_*i*_ = 1, the full conditional for each pair (*z*_*ij*_, *η*_*ij*_), holding all other amino acids fixed, is

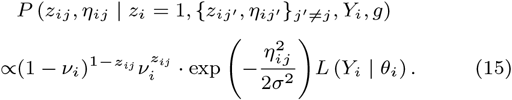

Since Ψ_*i*_ is a joint function of all 20 pairs (*z*_*ij*_, *η*_*ij*_)_*j∈A*_, (15) does not factor across *j* and admits no closed-form draw. We therefore update each *j* ∈ A in turn via an independence Metropolis step, proposing directly from the prior,

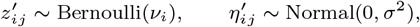

so that the prior terms cancel in the Metropolis ratio and the acceptance probability reduces to a likelihood ratio,

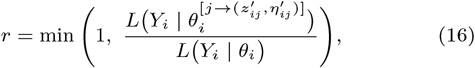

where 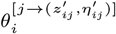 denotes *θ*_*i*_ with amino acid *j*’s components replaced by the proposed values. The likelihoods in (16) can be evaluated by pruning. When *z*_*i*_ = 0, Ψ_*i*_ does not depend on (*z*_*ij*_, *η*_*ij*_)_*j∈A*_ at all, and the conditional reduces to the prior, so each pair is drawn directly from its prior, independently across *j*, with no likelihood evaluation required.

Given the (updated) auxiliary parameters, the site inclusion indicator has full conditional

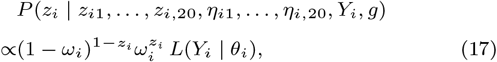

where *L*(*Y*_*i*_ | *θ*_*i*_) is evaluated at the current auxiliary values if *z*_*i*_ = 1, and at the uniform neutral profile if *z*_*i*_ = 0. Normalizing (17) yields the corresponding Bernoulli draw for *z*_*i*_.

### A.3 Gibbs Update for Fitness Profiles under Sufficient Augmentation

Under the sufficient augmentation, 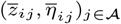 are instead defined conditionally on *z*_*i*_.

When *z*_*i*_ = 1, the full conditional combines prior and likelihood as

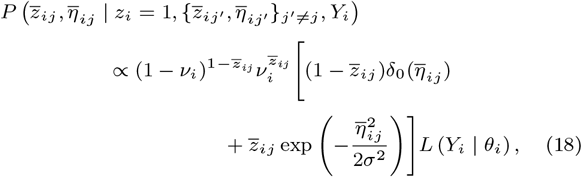

and is updated as in (15)–(16), by independence Metropolis sweeps over *j* ∈ A with proposals drawn from the prior. When *z*_*i*_ = 0, the hierarchy forces 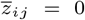 and 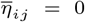 for every *j* ∈ A simultaneously, and this step is degenerate: no lower-level indicator or selection coefficient is retained whenever *z*_*i*_ = 0, so reactivating *z*_*i*_ requires the sampler to rebuild 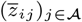 from scratch, one amino acid at a time. This explains the slow-mixing documented in Section 4.2.

Since 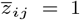 for any *j* implies *z*_*i*_ = 1, the site inclusion variable is 1 whenever 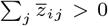. Otherwise, *z*_*i*_ is drawn from the conditional Bernoulli distribution

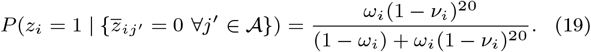

### A.4 Gibbs Update for Fitness Profiles with Interweaving Strategy

The accelerated sampler of Section 4.3 interweaves the ancillary and sufficient updates above within a single iteration at each site *i*. One iteration proceeds as follows.

i. Update 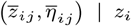 for each *j* ∈ A under the sufficient augmentation, as in (18).
ii. Update 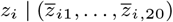 under the sufficient augmentation, as in (19).
iii. Map the updated values into the ancillary parameterization:
  - if *z*_*i*_ = 1, set 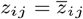 and 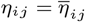 for all *j* ∈ A;
  - if *z*_*i*_ = 0, impute *z*_*ij*_ ∼ Bernoulli(*v*_*i*_) and *η*_*ij*_ ∼ Normal(0, *σ*^2^) directly from their priors.
iv. Update *z*_*i*_ | (*z*_*i*1_, … , *z*_*i*,20_, *η*_*i*1_, … , *η*_*i*,20_) under the ancillary augmentation, by proposing 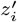 from its prior Bernoulli(*ω*_*i*_) and accepting with probability min(1, *ρ*), where

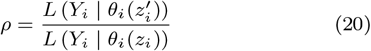

is the likelihood ratio between the rate matrices implied by 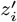 and by the current *z*_*i*_.

